# Impact of pre-breeding feeding practices on rabbit mammary gland development at mid-pregnancy

**DOI:** 10.1101/2022.01.17.476562

**Authors:** C Hue-Beauvais, K Bébin, R Robert, D Gardan-Salmon, M Maupin, N Brun, E Aujean, F Jaffrezic, S Simon, M Charlier, F Le Provost

## Abstract

Optimizing rabbit does preparation during early life to improve reproductive potential is a major challenge for breeders. Does selected for reproduction have specific nutritional needs, which may not be supplied with the common practice of feed restriction during rearing in commercial rabbit production. Nutrition during early life was already known to influence metabolism, reproduction and mammary gland development later in life, in particular during pregnancy. The aim of this study was to analyze the impact of four different feeding strategies in the early life of rabbit females (combination of high or moderate feed restriction from 5 to 9 weeks of age with restricted or ad libitum feeding regime from 9 to 12 weeks of constituting the pubertal period) on their growth, reproductive capacities and mammary development at mid-pregnancy.

Unlike food intake, which remains regular, mean body weight gain was inversely proportional to the dietary restriction applied over the considered periods. The feeding strategies in place for the four groups had no effect on the reproductive parameters of the females at mid-pregnancy, as opposed to certain metabolic parameters such as cholesterolemia, that decreased with dietary intake at puberty (p≤0.05). Furthermore, restriction programs have impacted mammary tissular structures at mid-pregnancy. The expression of lipid metabolism enzymes (Fatty acid synthase N and Stearoyl co-A desaturase) is also increased in mammary epithelial tissue at mid-pregnancy by the dietary strategies implemented (p≤0.05). Moreover, milk gene expression, used as differentiation markers, indicates a better mammary epithelial development regarding further lactation, in the case of the less restrictive strategies during early life period, especially the higher feeding allowance. Our results highlight the importance of investigating feeding conditions of young female rabbits and nutrition in early life rearing, in order to provide specific recommendations for optimizing lactation and thus preventing neonatal mortality of the offspring.

## INTRODUCTION

As a result, for the past 15 years, most rabbit breeders have implemented a feeding plan for growing rabbits (Gidenne et al., 2012). These strategies had the advantage of reducing the risk of post-weaning digestive disorders and improving feed efficiency in the animals (Gidenne et al., 2009), while for the breeder, they presented both economic and environmental impacts (Zened et al., 2013). As a result, in conventional rabbit farming in France, the cost of food represents 50% of the selling price of a rabbit (Cartuche et al., 2014).

In most breeding farms, post weaning (after 5 weeks of age) does are reared the same way as fattening animals (up to 10 weeks of age). Between 10 weeks of age and the first artificial insemination (AI) (19.5 weeks of age), young females receive a restricted quantity of feed daily. Nevertheless, feeding restriction, in particular during pregnancy, frequently causes energy deficit leading to poor fertility (Cappon et al., 2005; Matsuoka et al., 2006). Inadequate energy intake also impairs lactation and thus kit survival, growth and dietary transition, due to the level and quality of milk production (Hue-Beauvais et al., 2017).

Mammary gland is a complex secretory organ containing different tissues, one of the most important being mammary epithelial tissue, composed of different cell types, responsible for the synthesis and secretion of milk components. The growth and differentiation of mammary gland are long-term process that starts early in life and continues through adulthood, including reproductive cycles (Macias & Hinck, 2012). Early life factors, that may influence mammary development, can alter mammary development later at pregnancy and thus, impact the epithelial cell population, responsible for the synthesis and secretion of milk components during lactation (Robinson GW, 1995).

Nutrition influences mammary gland development, with an impact depending on critical periods of susceptibility, such as weaning, puberty, pregnancy or lactation (Hue-Beauvais et al., 2021). In species, such as rabbit, administration of an obesogenic diet, from the neonatal period or during puberty induces deleterious effects on metabolism and mammary gland development later in life (Olson et al., 2010; Hue-Beauvais et al., 2019; Hue-Beauvais et al., 2021). Rearing does with fibrous diets increased the ability of primiparous females to obtain resources, especially at the onset of lactation (Martinez-Paredes 2019).

Concerning the feeding strategies applied in breeding, studies showed detrimental effects on mammary gland development and metabolism induced by restriction followed by over allowance diet in gilts breeding (Farmer et al., 2012). In the same way, feeding restriction was associated with modification of mammary epithelial tissue and milk properties in cattle farming (Stumpf et al., 2013). Finally, the increase of the feeding level during the post-weaning rearing period could be an interesting way in goat breeding, to enhance body development without impairing mammary gland development whilst having a positive impact on reproductive parameters such as litter weight (Panzuti et al., 2019).

The effect of restricted feeding management over early stages of life, such as post-weaning and puberty on mammary gland development during pregnancy and subsequent lactation remains unknown in the rabbit. Although, some available literature does not support the hypothesis that feed restriction impairs the milk yield or quality (Martinez-Paredes et al., 2019), there is a lack on the knowledge on the impacts of a long-term feed restriction, starting as soon as 5 weeks old, on the physiological development of the mammary gland and tissues

In this study, we investigate the impact of four feed restriction strategies used in female rabbits breeding, over two distinct periods (post-weaning and puberty) on different physiological aspects such as growth, metabolic profiles, reproductive parameters and mammary gland development on day 14 of first pregnancy corresponding to the transition from the proliferative phase of the mammary cells to the differentiation of the lobulo-alveolar structures to form acini (Lu & Anderson, 1973).

## METHODS

### Animals, experimental design and sampling

This study was carried out in compliance with the French regulations on animal experimentation and with the authorization of the French Ministry of Agriculture. Protocol was approved by an Ethics Committee registered within the French Comité National de Réflexion Ethique sur l’Expérimentation Animale.

Forty female rabbits (Hyplus PS19) were housed individually in an indoor facility under controlled conditions of temperature (18°C) and light (usually an 8/16 h light/darkness cycle except for an inverted cycle during the week before mating). All rabbits received the same conventional commercial breeding diet (2,350 kcal of digestible energy, 15.2 % of crude protein, 17 % of celluloses, 0.56 % of digestible lysin). At weaning (5 weeks of age), the females were divided into two equivalent groups of 20 females, according to the body weight (862 ± 61 g). During the post-weaning period, from the fifth to the eighth week inclusive, the rabbits were weekly weighted and fed either with a strict restricted (SR; 7% of live weight at weaning + 2 g/d), or a moderate restricted (MR; 9% of live weight at weaning + 2,5 g/d) quantity of feed (Fig.1). During the pubertal period, for 3 weeks from 9 to 11 weeks of age included (Hulot et al., 1982), both groups were randomly divided to form four experimental groups of 10 females, which received diet at 140 g/d (groups SRR: SR-Restricted and MRR: MR-Restricted) or *ad libitum* (groups SRAL: SR-*Ad Libitum* and MRAL: MR-*Ad Libitum*) (Fig.1). At 12 weeks of age, rabbits were housed individually and received 150 g/d of diet. Then growth was determined by weighing the rabbits once a week to the age of 19 weeks. Food intake was also monitored on the same weekly basis during pubertal period for SRAL and MRAL groups by the difference between the weight of the distributed ration and the refusal. At 19 weeks of age, the females were mated by AI and pregnancy was confirmed by abdominal palpation on Day 14 of pregnancy. then euthanized, confirmed before euthanasia,The 40 pregnant females were euthanized by exsanguination at 14 days of pregnancy, after 12 hours of hydrous fasting allowing subsequent blood metabolic assays, to obtain mammary gland tissues samples.

**Figure 1:**
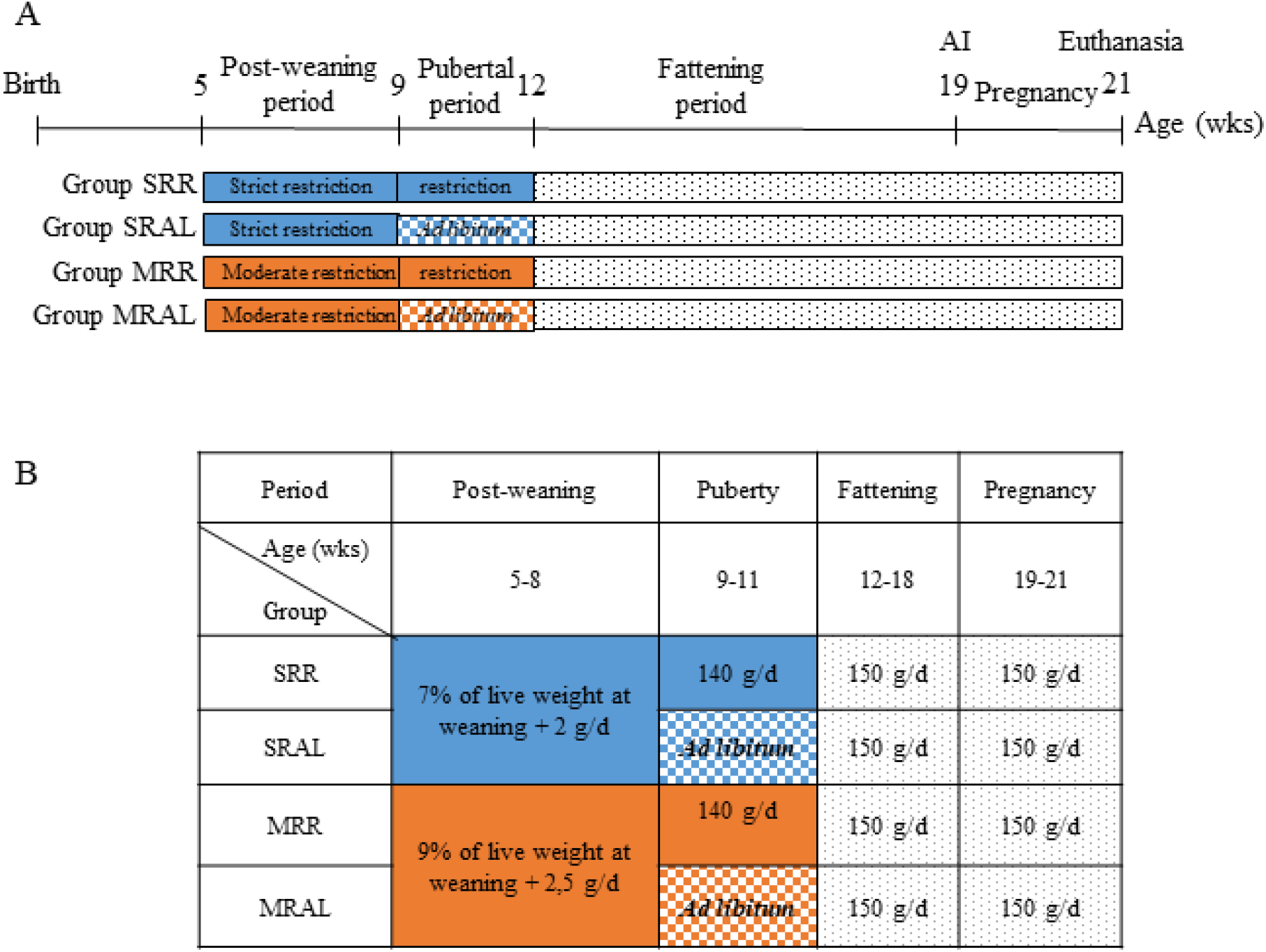
(A) Design of the experimental protocol. Each group (n = 10) received various quantities of food, over both post-weaning and pubertal periods. (B) Description of feeding strategies for each group of rabbits (SRR : strictly restricted during post-weaning period and restricted during puberty: SRAL : strictly restricted during post-weaning period and fed *Ad libitum* during puberty: SRR : moderately restricted during post-weaning period and restricted during puberty; MRAL. moderately restricted during post-weaning period and fed *Ad libitum* during puberty).

### Sampling and metabolic assays

Blood samples of each forty females were collected, at euthanasia by exsanguination, in tubes containing EDTA to determine levels of triglyceride (Triglyceride EnzyChrom kit; Cliniscience), cholesterol (Cholesterol RTU kit; Biomerieux), glucose (Glucose RTU kit; Biomerieux) and leptin (Cloud Clone Corp.). Left inguinal mammary gland from each 40 animal was fully excised and dissected to remove muscle, then mammary samples were processed and stored for further analyzes. All embryo vesicles were dissected, opened and the fetuses numbered and extracted to confirm viability by macroscopic examination.

### Histological analysis

For histology, samples were fixed for 24 hours in RCl2 buffer (Alphelys, France) before embedding in paraffin. Five-micrometer sections, separated at least by 100 μm each in the thickness of the tissue were mounted on slides. Slides were stained with hematoxylin and eosin (H&E; Sigma-Aldrich) and then digitized under bright light using a Hamamatsu NanoZoomer (Hamamatsu Photonics). Four sections per sample, i.e. eight sections per rabbit were processed and the percentage of areas occupied by mammary epithelial tissue (clusters of alveolar structures), adipose tissue, mammary duct lumens or connective tissue were measured using CaseViewer software (3D Histech) and divided by the whole section area to generate the proportion of each tissue. Results are expressed as means ± SEM.

### RNA Extraction and RT-qPCR analyses

Total RNA from mammary epithelial tissue was isolated from each mammary sample using the RNA NOW kit (Ozyme) according to the manufacturer’s protocol. The integrity of ribonucleic acid was assessed using an Agilent Bioanalyzer. Samples with an RNA Integrity Number (RIN) higher than 7 were subsequently used (Fleige & Pfaffl, 2006).

For quantitative PCR (qPCR) assays, reverse transcription (RT) was performed on 200 ng of each mammary sample’s total RNA using the SuperScript VILO cDNA Synthesis kit according to the manufacturer’s instructions (Invitrogen) and under the following conditions: 42°C for 60 min and 85°C for 5 min. qPCR runs were achieved using Applied Biosystems SYBR Green PCR Mastermix (Thermo Scientific) according to the manufacturer’s instructions, on a QuantStudio system (Thermo Scientific). After optimization of the qPCR systems (efficiency ranging from - 3.25 to -3.45), amplification reactions were run in triplicate under the following conditions: 95°C for 15 min, 45 cycles of 95°C for 15 sec and 60°C for 1 min. The threshold cycles obtained for each gene were normalized with the values of the *TATA Binding Protein (Tbp)* gene and the results were expressed as fold changes of the threshold cycle (Ct) values relative to the control using the 2-ΔΔCt method (Livak & Schmittgen, 2001). The primers used for each gene (*κ-casein, Whey acidic protein, α-lactalbumin, Fatty acid synthase N and Stearoyl co-A desaturase*) amplified are presented in Table 1.

**Table 1:**
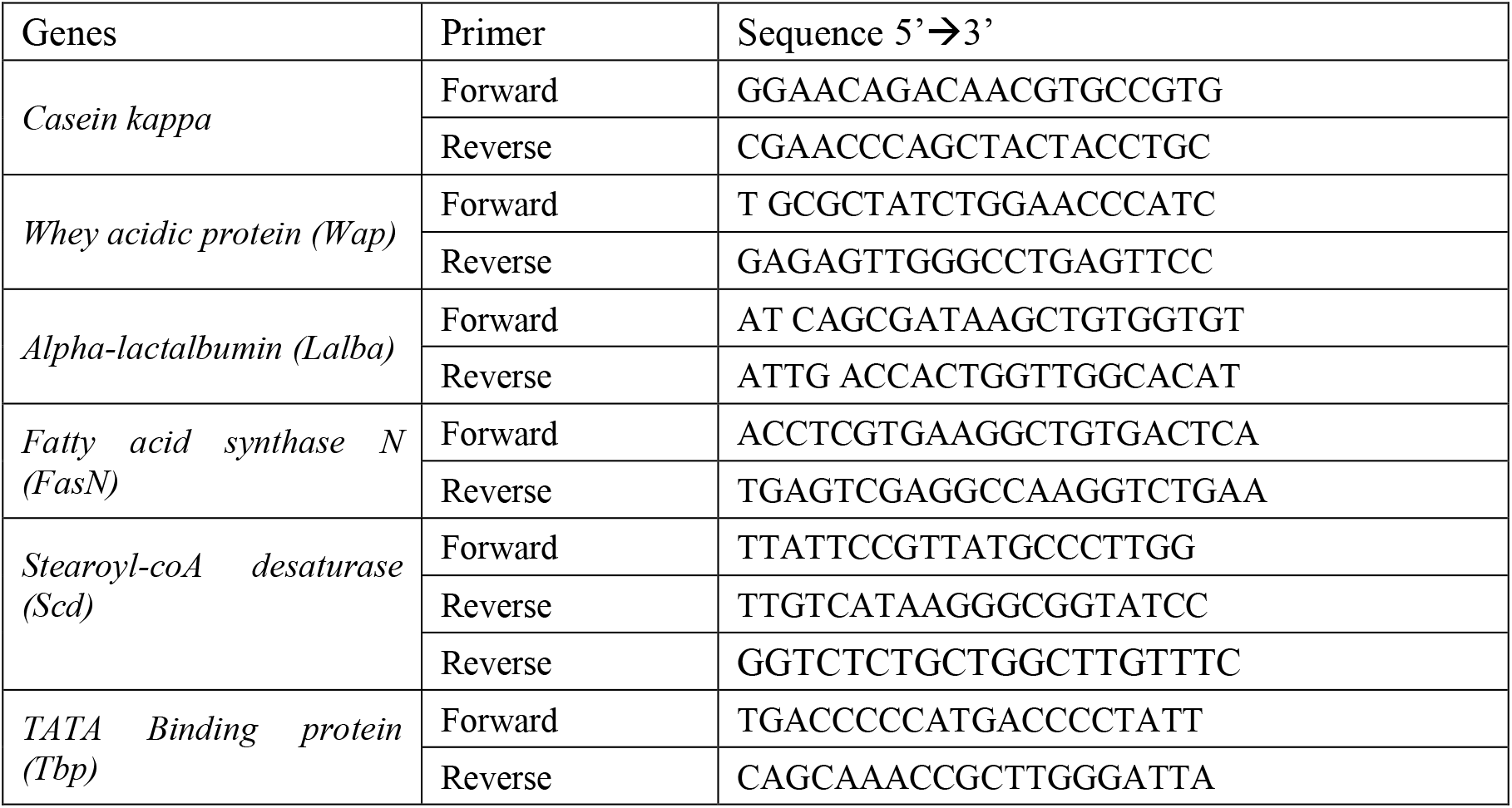
Primer sequences used for qPCR experiments

### Statistical analyses

Concerning rabbit growth, two periods were analyzed separately: 1) the post-weaning and pubertal period, 2) the fattening period. In order to take into account the longitudinal nature of the data, a linear mixed model was used for both periods using the nlme R package, with a random animal effect and a first order autoregressive correlation structure. For the fixed effects, a linear curve was first considered, with the time effect as continuous. The fixed effects were therefore in this case the group effect, the continuous time and an interaction between group and time.

In order to have a more precise analysis of the differences among groups between times we performed a second type of analysis with the time effect as discrete (per week), and the lsmeans option to specify the contrasts of interest. Once again, the slopes over time were found to be significantly different between groups, as well as the intercept at the beginning of the fattening period.

Experimental data are presented as means±SEM (standard error of the mean). Statistical analyses were performed to detect significant inter-group differences using either unpaired Student’s *t*-test when the sample size was >30, or the Mann-Whitney U-test when the sample size was <30). *P* ≤ 0.05 was considered to be significantly different.

## RESULTS

Different levels of restriction during 2 separate periods, covering the early-life from 5 to 12 weeks of age have been compared (Fig. 1). During the first period of restriction which covers the post - weaning period and lasts for 4 weeks, one group (SR group) received a more drastic restriction than the second group (MR group) (quantity of feed: SR group 95 g/d and MR group 120,75 g/d). At the beginning of the post-weaning period, mean body weight of does was 832±61 g, equally distributed among SR and MR groups. It was observed for the first period that the differences between groups at the beginning of the period were not significant. The differences between slopes over time were however significantly different between groups. During the first three weeks, from 5 to 7 weeks of age, body weight was similar between group (Fig 2), but from 8 weeks of age, does of group MRR weighted higher than those of group SR (P=0.015).

**Figure 2:**
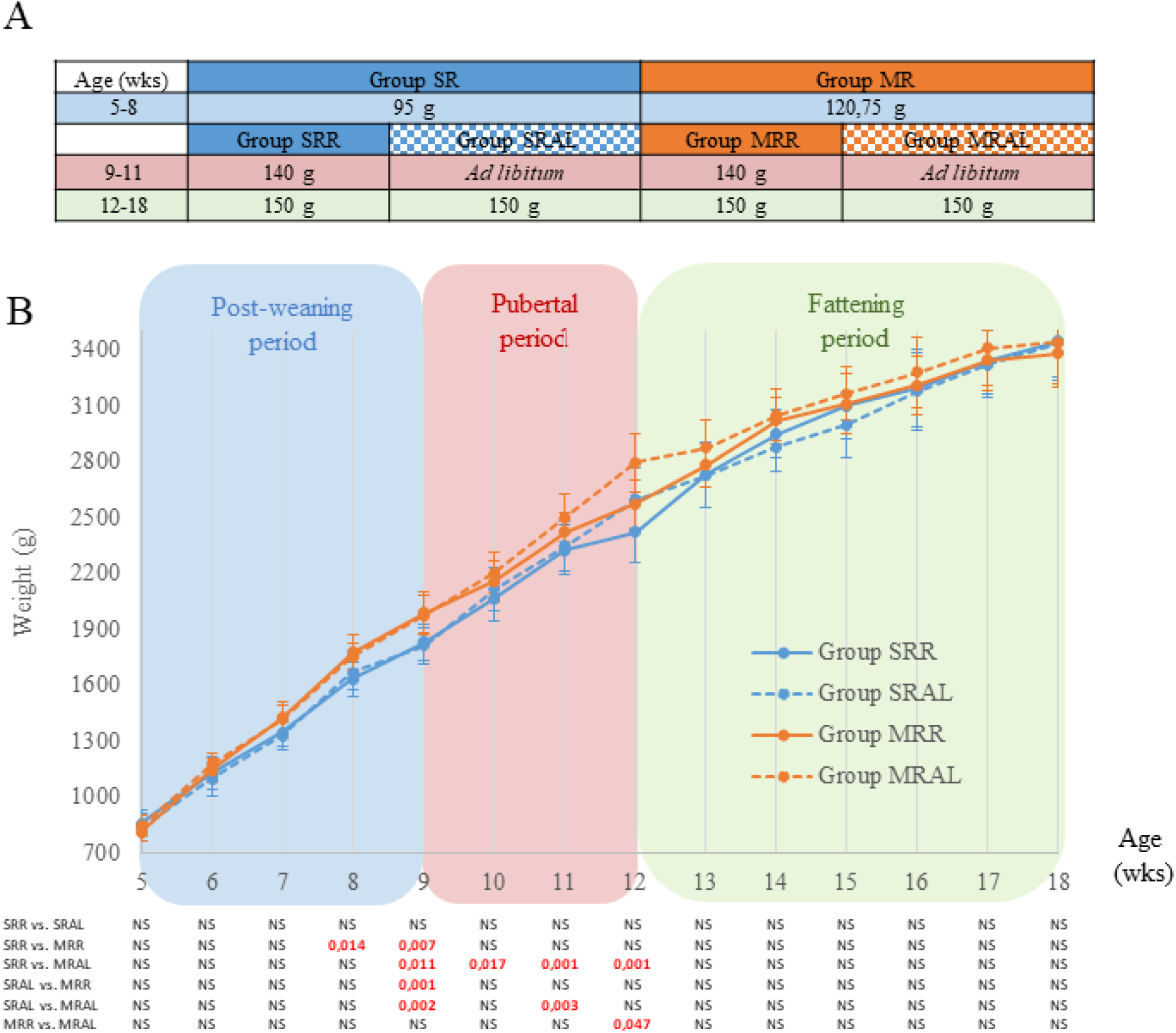
Effect of the four feeding strategies over three periods of age. in female rabbits. (A) Average daily food intake. Data are expressed by Mean ± sem. (: SRR : strictly restricted during post-weaning period and restricted during puberty; SRAL : strictly restricted during post-weaning period and fed *Ad libitum* during puberty; SRR: moderately restricted during post weaning period and restricted during puberty; MRAL : moderately restricted during post-weaning period and fed *Ad libitum* during puberty) (B) Body weight, weekly measured in the four different groups of female rabbits, between 5 and 18 weeks of age. Data are expressed by Mean ± sem. Significant differences (P < .05) between the groups are indicated below in red. NS means Non Significant.

At 9 weeks of age, each group was divided into 2 sub-groups according to the quantity of food received, restriction feeding with 140 g/d (groups SRR and MRR) or *ad libitum* (groups SRAL and MRAL).

The difference of weight observed during the late post-weaning period remained significant during the pubertal and fattening periods and depends on the restriction feeding pattern. At the end of the pubertal period, from 9 to 11 weeks of age, group SRR rabbits have the lowest weight curve (P≤0.01). At 12 weeks of age, group MRAL animals show a higher body weight than group SRR rabbits (2795± 155g and 2421±99g respectively, P<0.001), and the same difference is observed between MRR and MRAL groups (2571±130g and 2795± 155g respectively, P=0.047): rabbits strongly restricted in post weaning and fed *ad libitum* during puberty (group SRAL), reached the same weight than rabbits less restricted in post-weaning and restricted during pubertal (group MRR). Furthermore, whatever the diet during the post-weaning period, animals fed *ad libitum* during the puberty period are heavier than those fed restricted diets (MRAL mean weight higher than SRR and MRR Fig 2B). As the groups SRAL and MRAL were fed *ad libitum*, the mean food intake was calculated during the pubertal period. No significant difference was observed between both groups.

Once again, the slopes over time were found to be significantly different between groups, as well as the intercept at the beginning of the fattening period. (12 weeks of age). The weight of the does was higher in the less restricted group (MRAL) than in the SRR and MRR groups (2795±155 g for MRAL, 2597±167 g for SRAL and 2571±130 g for MRR, p<0.05) which were similar to each other, and in the SRR group in which the mean weight was the lowest (2421±99 g, p>0.05). From the 13^th^ week of age until the end of the fattening period (18^th^ week of age) during which all the rabbits were receiving the same quantity of feed, body weights of groups were not significatively different. At the end of the fattening period, body weights were equivalent within the four groups (Fig 2B).

Metabolic profiles were evaluated at mid-pregnancy by measuring glucose, cholesterol, triglycerides and leptin in blood after 12 hours of fasting (Fig 3). No difference was observed between the four feeding strategies concerning glycemia, triglyceridemia or leptinemia at mid-pregnancy. Unrestricted feeding during pubertal period when animals received a restricted feed during post-weaning period (groups SRR vs. SRAL) provoked a significant decrease in cholesterol levels (0.25±0.02 g/L for SRR group and 0.19± 0.02 g/L for SRAL group, p=0.045, Fig 3). This difference was not observed among does which were less restricted during the post-weaning period (groups MRR vs. MRAL).

**Figure 3:**
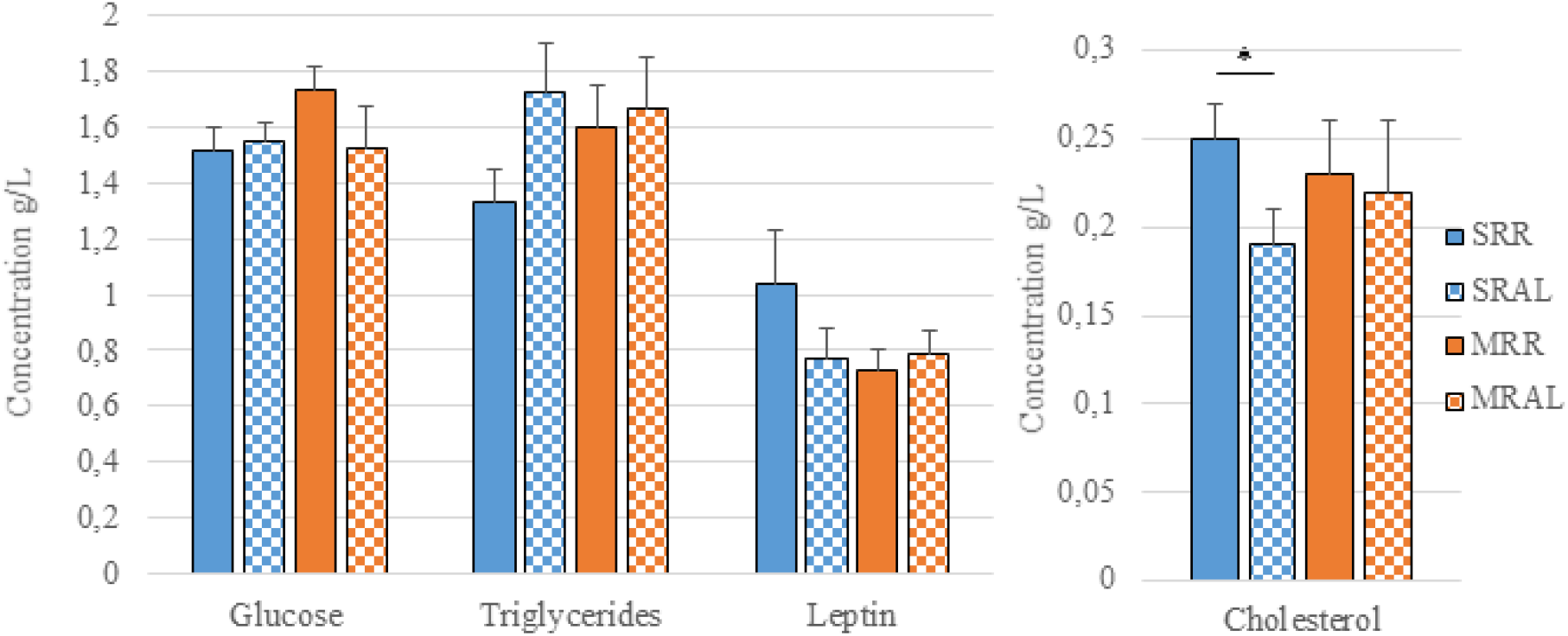
Metabolic profiles of each group in (= 10) at mid-pregnancy. : SRR : strictly restricted during post-weaning period and restricted during puberty; SRAL : strictly restricted during post-weaning period and fed *Ad libitum* during puberty; SRR : moderately restricted during post-weaning period and restricted during puberty; MRAL : moderately restricted during post-weaning period and fed *Ad libitum* during puberty; Data are expressed as means ± SEM. Significant differences (p <0.05) between the groups are indicated by asterisks (*).

The feeding strategies had no effect on the reproductive parameters: the fprolificacy a nd the fetal viability (Fig 4) were not different in the four groups.

**Figure 4:**
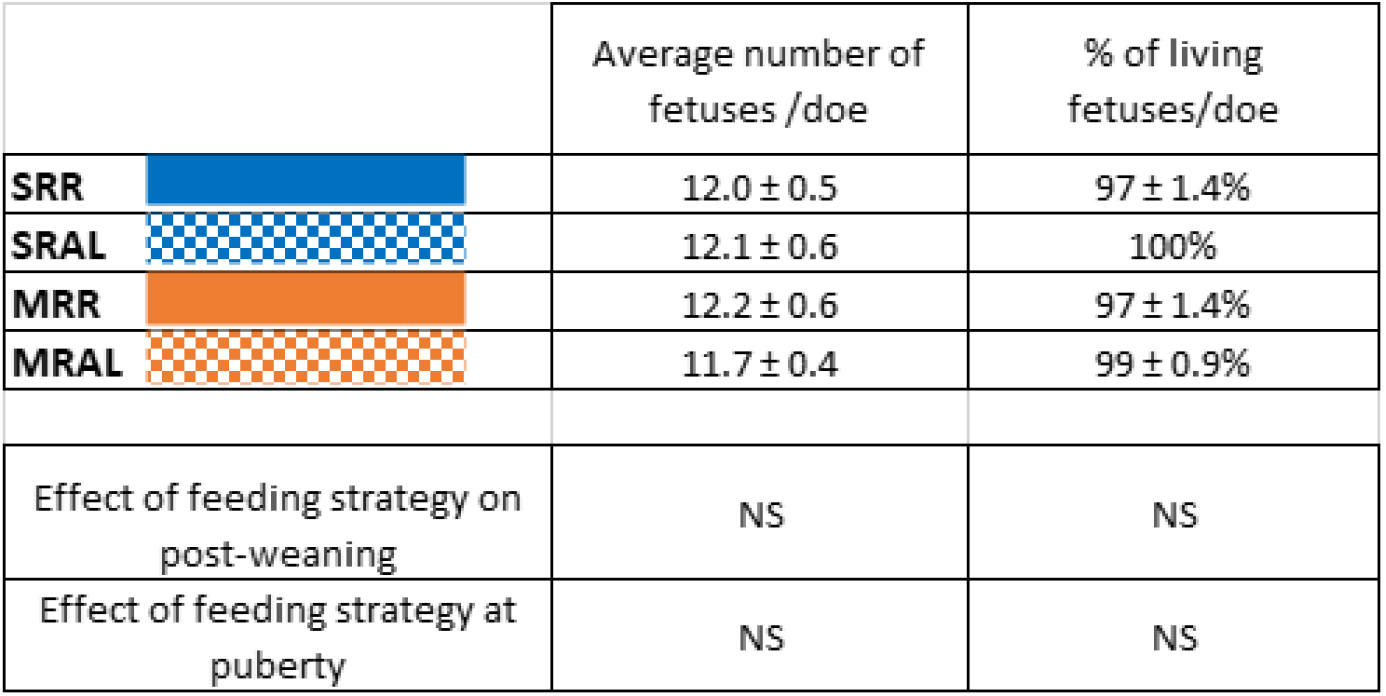
Average number of fetuses and average percentage of living fetuses per doe in each group (n = 10). SRR : strictly restricted during post-weaning period and restricted during puberty; SRAL : strictly restricted during post-weaning period and fed *Ad libitum* during puberty; SRR : moderately restricted during post-weaning period and restricted during puberty; MRAL: moderately restricted during post-weaning period and fed *Ad libitum* during puberty. Mean ± sem. NS means Non Significant.

To examine the effects of the feeding strategies during both post-weaning and pubertal periods on mammary gland, histological analyses were performed. The examination of mammary tissue sections from all does on Day 14 of pregnancy revealed some differences between the four groups (Fig 5). The surface occupied by mammary connective tissue increased, while adipose and epithelial tissues decreased in group SRAL compared to group SRR (Fig 5B). This difference was not observed when does received a less restricted diet during the post-weaning period (groups MRR vs. MRAL). The area corresponding to the ducts’ lumina was not significantly different within the four groups. (Fig 5B), although a declining trend can be observed in the less-restricted groups during the post-weaning period (MRR and MRAL compared to SRR or SRAL groups; p=0.06 and p=0.056, respectively).

**Figure 5:**
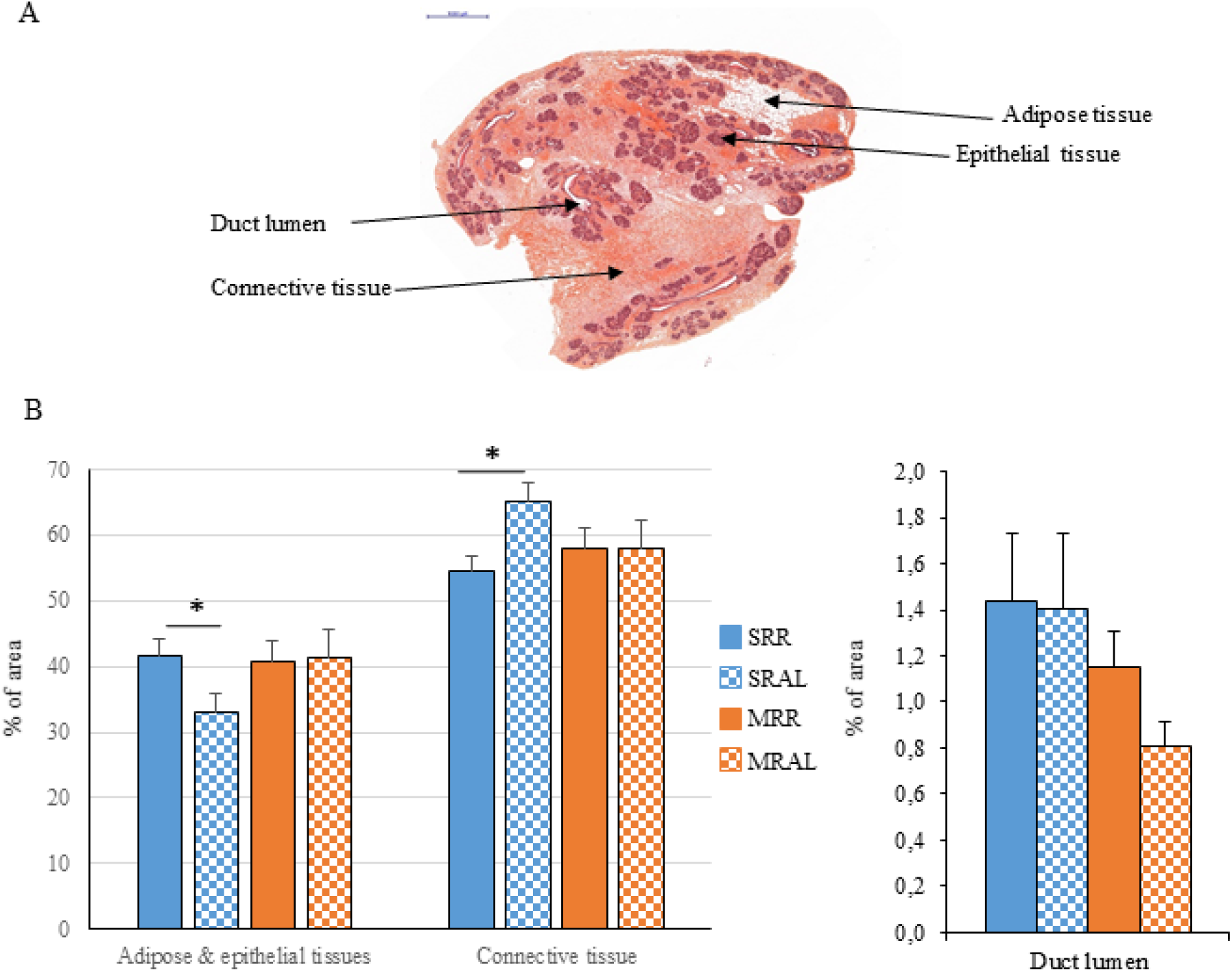
Histological analyses of mammary gland on Day 14 of pregnancy in each group (SRR : strictly restricted during post-weaning period and restricted during puberty; SRAL : strictly restricted during post-weaning period and fed *Ad libitum* during puberty; SRR : moderately restricted during post-weaning period and restricted during puberty. MRAL : moderately restricted during post-weaning period and fed *Ad libitum* during puberty) (A) Representation of the different types of tissues present on a representative histological section of mammary gland, stained with hematoxylin and eosin. Scale bar represents 1mm (B) Relative quantification of areas occupied by the different types of mammary tissue: adipose and epithelial tissues, connective tissue: and duct lumen. Data are expressed as means ± SEM. Significant differences (P < 0.05) between the groups are indicated by asterisks (*). Number of animals per group n = 10.

In order to study the modifications induced by the different feeding strategies, mammary epithelial cell differentiation status, using mammary differentiation markers such as enzymes involved in lipid metabolism (*Fatty acid synthase N (FasN)* and *Stearoyl-coA desaturase* (*Scd*)) as well as milk proteins (*kappa casein, Whey acidic protein* (*Wap*) and *alpha-lactalbumin* (*Lalba*)) were assessed. *FasN* expression increased when less restricted feeding strategies were used (Fig 6).

**Figure 6:**
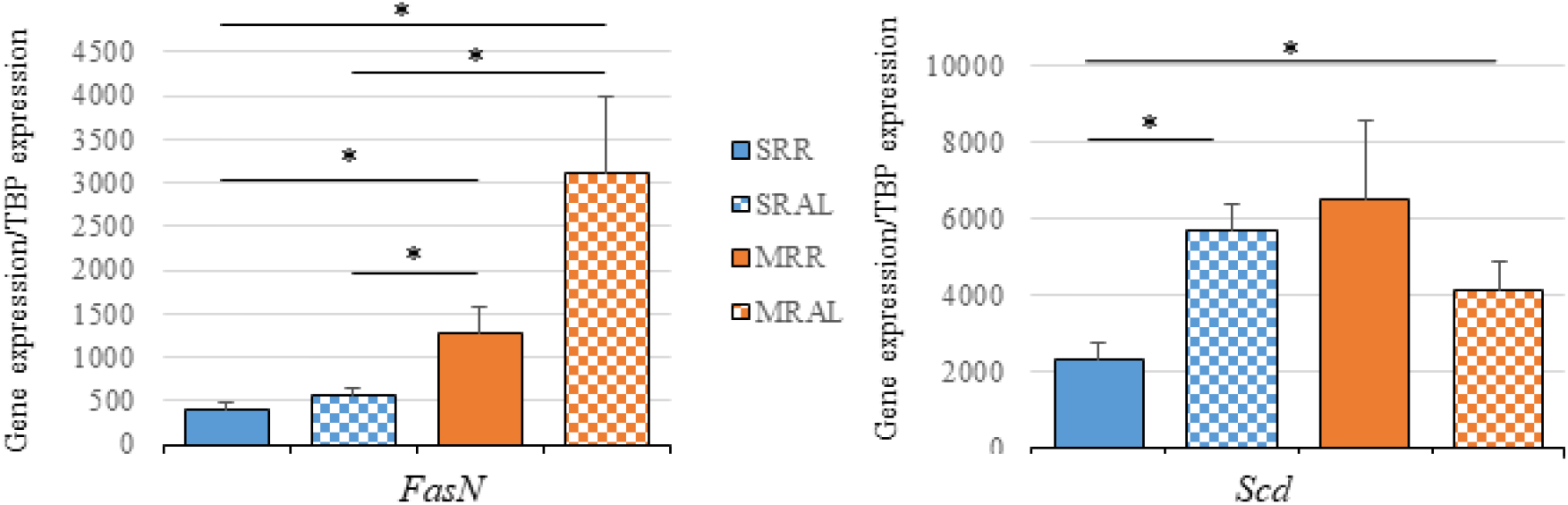
Expression of genes involved in lipid metabolism in mammal epithelial tissue on Day 14 of pregnancy. Analyses were performed in the four groups of rabbits (n = 10 in each group) according to the feeding strategy (SRR : strictly restricted during post-weaning period and restricted during puberty; SRAL : strictly restricted during post weaning period and fed *Ad libitum* during puberty; SRR: moderately restricted during post-weaning period and restricted during puberty; MRAL: moderately restricted during post weaning period and fed *Ad libitum* during puberty) The expression of transcripts was assessed for *Scd* and *FasN* and normalized with *Tbp* as housekeeping gene. Data are expressed as means ± SEM. Significant differences (at least *P* <0.05) between the groups are indicated by asterisks (*).

Indeed, higher levels of *FasN* transcripts were observed in groups MRR and MRAL than in groups SRR and SRAL, and these differences were strongly significant. *Scd* transcript levels increased between groups SRR and SRAL and MRAL, but were similar in groups SRAL, MRR and MRAL (Fig 6).

To further investigate the changes induced by the different feeding strategies in mammary epithelial tissue, we analyzed the patterns of expression of genes encoding milk proteins (*kappa casein, Whey acidic protein* (*Wap*) and *alpha-lactalbumine* (*Lalba*)), using RT-qPCR (Fig. 7), since these genes are specific markers for differentiated mammary epithelial cells (MEC). In all groups a high individual variability within the MRR group was observed. However, analyses revealed no difference between groups concerning the □-*casein* gene expression. *Wap* expression profile was similar between SRR and SRAL as well as MRR and MRAL groups, The *Lalba* transcript level was higher in MRAL than in SRAL group (P=0,041) highlighting an effect of feeding strategyduring post-weaning period on the expression of this gene when the animals are fed *ad libitum* during pubertal period.

**Figure 7:**
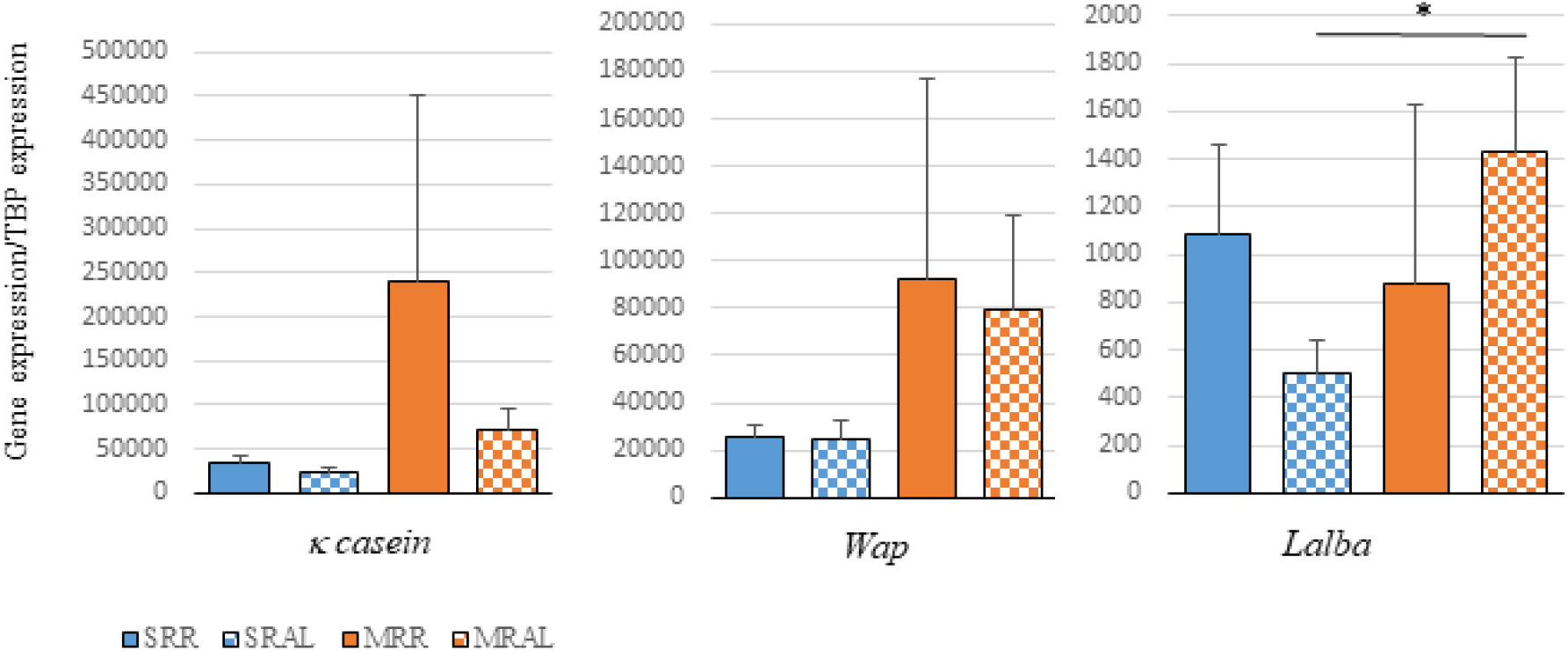
*K casein, Wap* and *Lalba* expression in mammary epithelial tissue on day 14 of pregnancy of the four groups of rabbits (n = 10 in each group), according to the feeding strategies (SRR : strictly restricted during post-weaning period and restricted during puberty; SRAL : strictly restricted during post-weaning period and fed *Ad libitum* during puberty; SRR : moderately restricted during post-weaning period and restricted during puberty; MRAL : moderately restricted during post-weaning period and fed *Ad libitum* during puberty). Transcripts levels were assessed for *κ casein, Wap* and *Lalba*, and normalized with *Tbp* as housekeeping gene. Data are expressed as means ± SEM. Significant differences (at least *P* <0.05) between the groups are indicated by asterisks (*).

## DISCUSSION

For economic and logistical reasons, does, whether they are intended for meat production or to become the future breeders, are often fed the same way in the farms. This means that the animals are subjected to restrictive feeding strategies during their early-life periods, such as post-weaning and puberty. Here, influence of different combinations of dietary restrictions was studied, with a particular focus on metabolism, number and viability of fetuses and mammary gland development, in rabbits at mid-pregnancy.

Our results showed that restrictive feeding changes starting at the post-weaning period provoke a difference in body weight. This difference occurs from the third week of diet, and is due to a difference of approximately 25g/d in the quantity of feed, thus emphasizing the high sensitivity of growing rabbit to nutrition during the post-weaning period. Such effects have already been observed in rabbits submitted to a short intensive food restriction followed by a re-alimentation period (Tumova et al., 2016). Feeding strategies during pubertal period have shown that *ad libitum* groups (SRAL and MRAL) have increased food intakes (> 150 g/d) than restricted groups SRR and MRR, which received 140 g/d. As expected consequently, body weight is higher in the less restricted-fed groups (SRAL and MRAL) at the end of pubertal period. These results underline the importance of considering young life, including post-weaning and puberty, as a critical period for nutrition, and where feed restriction could alter normal development. Switching during the fattening period to a quantitatively identical diet for all four groups harmonized the body weight of all the does. Moreover this is consistent with studies showing that irregular weight growth in response to feeding has long-term consequences in adulthood (Haschke et al., 2019; Neave et al., 2019).

Nevertheless, mechanisms involved still remain unclear in particular regarding the impact of feeding restriction on reproduction (Villeneuve et al., 2010b). According to previous work, restricted feeding strategies tested did not impair reproductive performances at pregnancy, while it has been showed, in ovine that specific plan of food restriction could affect the onset of puberty, and negatively influence the hypothalamic-pituitary-ovary axis involved in the hormonal control during pregnancy (Villeneuve et al., 2010a ; Wang et al., 2016; Rizzoto et al., 2019).

Measurements of blood metabolic parameters are commonly used to assess the effect of diets on body physiology. As already showed in poultry (Yang et al., 2010) analysis of metabolic parameters at mid-gestation in does showed a higher level of plasma cholesterol in the most restricted group showing that feed restriction in post-weaning produce long-term effect on lipid metabolism. This finding could be also related to stress induced by restriction feeding since a relationship has been observed between plasma cholesterol and leptin concentrations and stress-induced disorders (Jow et al., 2006; Shankar et al., 2012). Nevertheless, in our studies we only observed subtle differences when metabolic parameters were measured 9 weeks after the end of the feeding strategies. This may be explained by the fact that applied restriction strategies were weak.

In early life, the mammary gland is a potent target for environment effects, particularly those related to nutrition (Hue-Beauvais et al., 2019), because the mammary epithelium has entered a stage of growth (Denamur, 1963; Borellini & Oka, 1989) leading to the establishment of structures that will differentiate to produce milk components (Borellini and Oka 1989). The first half of gestation is essentially dedicated to the proliferation of mammary epithelial cells (MEC), while the second half is rather characterized by the differentiation of these cells. At mid-pregnancy (Day 14), interstitial adipose tissue gradually disappears and proliferating MEC fill in the inter-ductal spaces and start to express genes that can be considered as differentiation markers, such as milk protein genes (Robinson 1995). This is a particularly opportune time to estimate and analyze the mammary consequences of nutritional changes that occurred in early life. Moreover, changes to mammary gland development during pregnancy can impact the MEC population that is responsible for the synthesis and secretion of milk components in lactation (Robinson 1995).

No drastic changes in mammary gland histology were observed according to the different feeding strategies, which is reassuring since these strategies are based on restrictions practiced in the farms. However, a significant increase in connective tissue was observed in group SRAL compared to SRR, due to a combined decrease in fat and epithelial tissues. Within the mammary ducts, an tend to increased luminal area in the most restricted animals during the post-weaning period compared with the less restricted animals was observed. In addition, the weaker ductal areas were measured in not restricted rabbits during puberty.

Our results suggest that a severe feeding restriction, during post-weaning period, followed by *ad libitum* feeding during puberty may have deleterious effects on the mammary gland development and disturb its tissular composition, as observed here in pregnancy.

Consistent results were found in gilts showing that the impact of a strong restriction followed by unregulated feeding had negative consequences, among others on mammary gland development (Farmer et al., 2004; Farmer et al., 2012). This dietary dichotomy between these two life stages appears to be more deleterious than a restriction throughout the only post-weaning period. Indeed, in case of prolonged restriction, the body adapts and allows the preservation of mammary gland development (Park et al., 1994).

Lipid metabolism in mammary tissue was examined, using characteristic enzyme expression to correlate the defect in epithelial and adipose development with putative modifications in mammary function and differentiation. Fatty acid synthase N (FASN) and Stearoyl-coA desaturase (SCD) are both involved in the fatty acids biosynthesis and are found in cell types with a high lipid metabolism, such as MEC (Suburu et al., 2014). FASN is a rate-limited enzyme for fatty acid *de novo* biosynthesis in particular the long-chain fatty acids biosynthesis (Shi et al., 2015). *Fatty acid synthase N (FasN) gene expression was* increased in mammary glands of does with less restrictive nutritional status (SRR<SRAL<MRR<MRAL), which is consistent with the fact that intensive lipogenesis is correlated with higher level of nutrient intakes (Takeuchi et al., 2001).

Stearoyl-CoA desaturase (SCD) regulates membrane fluidity by the conversion of endogenous and exogenous saturated fatty acids into mono-unsaturated fatty acids (Angelucci et al., 2018). Overall, *Scd* gene expression appeared to decrease with high dietary restriction. Interestingly, increased levels of *Scd* transcripts were found in the group with the lowest cholesterol level (SRAL), thus contributing to correlate high SCD concentrations to metabolic disorders (Igal, 2011; Tsiplakou et al., 2015) as well as confirming the inversely proportional relation between cholesterol and SCD (Tian et al., 2018).

To determine whether feeding strategies during post-weaning and/or pubertal periods can affect MEC differentiation, expression of milk protein genes was performed. Kappa casein, WAP and α-Lactalbumin, are expressed exclusively in the MEC and can be used as MEC differentiation markers.

The three milk proteins’ transcripts tend to be increased in the less restriction feeding groups during post-weaning period (SRAL group *vs*. MRAL group), suggesting MEC differentiation. Those results in milk gene expression might indicate a better commitment toward mammary epithelial tissue function and lactation with the less restrictive strategies, especially with higher feeding allowance in early life period.

The impact of diet and nutritional strategies has been studied in different species, especially in farm animals. Among these, mammary gland development and the resulting milk production capacity has been related mainly to the qualitative aspect of the diet (hypo- or hypercaloric diet, low protein diet, etc.) (Fernandez-Twinn et al., 2010; Hue-Beauvais et al., 2011; Bautista et al., 2013). The impact of moderate feeding restriction on mammary gland development remains a little studied subject, depends on the species considered, and on the fate of the animal whether it is raised for meat, milk production or reproductive capacity. In ovine, effects of restricted feeding before puberty did not shown any negative impacts on mammary gland development, reproduction, lactation and offspring growth performance (Villeneuve et al., 2010a, 2010b). In the case of pig farming, the balance between feeding and mammary gland development is much more delicate (Farmer, 2018).

In rabbit breeding, feed restriction strategy is widespread used to improve productivity, decreased mortality and economic cost. For fattening rabbits, feed restriction during rearing period allowed uniformity in body weight and decreased neonatal mortality (Rommers et al., 2001). While a moderate feeding restriction may improve some sperm morphologic characteristics, as well as fertility in male rabbits (Pascual et al., 2016), it seems that restriction strategies have shown less beneficial effects and even more for rabbit does (Birolo et al., 2020).

Restricting feeding during different stages of pregnancy, even if it does not strongly affect growth of young rabbits, may delay placental growth, decrease the offspring survival and birth weight (Rommers et al., 2004; Manal et al., 2010 ; Matsuoka et al., 2012). In early life, rearing does with the same feeding strategies as fattening rabbits, leads to energy deficit, body mobilization and may reduce reproductive performance (Fortun-Lamothe, 2006). Consequently, feeding strategies and lifespan are closely linked in rabbit does.

Our findings showed that feed restriction strategies applied during post-weaning and/or pubertal period can impact mammary gland structure and may delay mammary epithelial tissue development and functionality. These results also suggest the urgent need of further investigations on milk composition and subsequent lactation capacity of restricted does through several reproductive cycles to provide recommendations. Indeed, while a moderate feeding restriction does not necessarily have consequences, a severe one could adversely affect health and breeding performances.

## Data accessibility

Dataset V3 are available online https://doi.org/10.5281/zenodo.6786934

## Acknowledgements

Version 3 of this preprint has been peer-reviewed and recommended by Peer Community In Animal Science (https://doi.org/10.24072/pci.animsci.100013).

